# Corticospinal excitability is facilitated during coordinative action observation and motor imagery of adapted single-leg sit-to-stand movements in young healthy adults

**DOI:** 10.1101/2024.12.11.627678

**Authors:** Ashika Chembila Valappil, Neža Grilc, Federico Castelli, Samantha Chye, David J. Wright, Christopher J. Tyler, Ryan Knight, Omar S. Mian, Neale A. Tillin, Adam M. Bruton

## Abstract

Combined action observation and motor imagery (AOMI) facilitates corticospinal excitability (CSE). This study used single-pulse transcranial magnetic stimulation (TMS) to explore changes in CSE for *coordinative* AOMI of a single-leg sit-to-stand (SL-STS) movement. Twenty-one healthy adults completed two testing sessions, where they engaged with baseline (BL), action observation (AO), and motor imagery (MI) control conditions, and three experimental conditions where they observed a *slow-paced* SL-STS while simultaneously imagining a *slow-* (AOMI_HICO_), *medium-* (AOMI_MOCO_), or *fast-paced* (AOMI_LOCO_) SL-STS. A TMS pulse was delivered to the right leg representation of the left primary motor cortex at three stimulation timepoints aligned with peak EMG activity of the knee extensor muscle group for the *slow-paced* (T3), *medium-paced* (T2), and *fast-paced* (T1) SL-STS during each condition. Motor evoked potential (MEP) amplitudes were recorded from the knee extensor muscle group as a marker of CSE for all stimulation timepoints and conditions. A main effect for experimental condition was reported for all stimulation timepoints. MEP amplitudes were significantly greater for AOMI_HICO_ at T1 and T3, and AOMI_MOCO_ and AOMI_LOCO_ at all stimulation timepoints, when compared with control conditions. This study provides neurophysiological evidence supporting the use of *coordinative* AOMI as an alternative method for movement (re)learning.

## Introduction

Action observation (AO) is the process of purposefully observing human movement via video or live demonstration (Eaves et al., 2022), while motor imagery (MI) involves the internal mental rehearsal of one’s own movement execution (MacIntyre et al., 2013). Both AO and MI are effective interventions for enhancing movement execution and (re)learning in rehabilitation and care settings (e.g., Buccino, 2014; Herranz-Gómez et al., 2020; Nicholson et al., 2019). These improvements have been explained by motor simulation theory, which posits that AO and MI activate brain areas associated with movement planning, production, and control, and thus have the capacity to increase functional connectivity across these motor networks (Jeannerod, 2001). In support of this proposition, research employing a range of neuroscientific modalities (i.e., fMRI, fNIRS, EEG, TMS) demonstrates increased cortico-motor activity for both AO and MI compared to control/baseline conditions (see e.g., Condy et al., 2021; Grosprêtre et al., 2015; Hardwick et al., 2018). Literature to-date has typically compared these two forms of motor simulation, showing that brain activation patterns for AO and MI are distinct, but overlapping (Hardwick et al., 2018), and that both AO and MI facilitate corticospinal excitability (CSE) as a marker of corticomotor activity compared to control conditions (e.g., Clark et al., 2004; Williams et al., 2012). After an initial review by Vogt et al. (2013), a body of work has investigated the behavioral outcomes and underlying neurophysiological mechanisms associated with the combined and simultaneous use of AO and MI.

Combined action observation and motor imagery (AOMI) involves a person observing human movement and simultaneously imagining the physiological sensations and kinesthetic experiences associated with executing the same or a different movement (Eaves, Riach, et al., 2016, Eaves et al., 2022). A recent meta-analysis by Chye et al. (2022) synthesized literature on highly-coordinated AOMI (AOMIHICO), where a person watches a movement demonstration and imagines the feeling of executing an identical movement in the same perspective in time with the viewed content. This meta analysis reported improved movement outcomes and facilitation of CSE for AOMI compared to AO and control conditions, but not compared to MI conditions. The behavioral and neurophysiological findings from Chye et al.’s (2022) meta-analysis suggest that AOMI_HICO_ benefits motor execution due to strong and widespread activation of movement-related brain regions. Specifically, repeated engagement in AOMI_HICO_ is likely to induce plastic-like changes in the motor system, benefitting physical execution of the movement in a similar manner to physical training (Chye et al., 2022; Grilc et al., 2024). This sentiment is supported in a rehabilitation context, as studies employing TMS show that AOMI_HICO_ leads to increased facilitation of CSE for whole-body functional movements such as walking (e.g., Kaneko et al., 2018) and balance (e.g., Mouthon et al., 2015, 2016), and studies employing AOMI_HICO_ as a training intervention report improved muscle strength (Scott et al., 2018), rehabilitation outcomes (Marusic et al., 2018), and postural control (Taube et al., 2014) across the lifespan.

Whilst empirical support for AOMI_HICO_ is growing, there is limited research on alternative forms of AOMI. Vogt et al. (2013) proposed that AOMI can occur along a spectrum, whereby AO and MI serve different roles during AOMI depending on the level of coordination between the two. At one end of this spectrum is *congruent* AOMI, where the content and/or visual perspective are identical for AO and MI components of AOMI. The term AOMI_HICO_ has been adopted instead of *congruent* AOMI in this study, as it has been proposed that the AO and MI components are never fully *congruent* during AOMI (Frank et al., 2020). This proposition was largely based on the different modalities adopted for AO (visual and auditory) and MI (kinesthetic) components in Vogt et al.’s (2013) definition for *congruent* AOMI, however it is worth noting that parameters such as body structure, movement pattern, and imagery ability will also result in an offset between AO and MI during AOMI of the same action, making AOMI_HICO_ a more suitable term. On the other end of the spectrum lies *conflicting* AOMI, where the content and/or visual perspective for AO and MI components of AOMI are opposing (e.g., watching movement and imagining being still and relaxed). In between these endpoints lies *coordinative* AOMI, where the content and/or visual perspective for AO and MI components of AOMI is similar. For this type of AOMI, the level of coordination between the observed or imagined actions can vary on a range of movement parameters, including the type, direction, speed, force, and accuracy of the movement (Chye et al., 2022; Grilc et al., 2024).

Two main theoretical hypotheses have been proposed to explain the underlying mechanisms for *coordinative* AOMI. The first is the Dual-Action Simulation Hypothesis (DASH) account for AOMI proposed by Eaves, Riach et al. (2016). Drawing from the Affordance-Competition Hypothesis (Cisek, 2007), the DASH suggests that individuals generate separate motor representations for the observed and imagined actions during AOMI, and that these are maintained as parallel sensorimotor streams that either merge or compete depending on the level of coordination between the observed and imagined actions. For AOMI_HICO_ these likely merge as one sensorimotor stream, producing more widespread activation in the premotor cortex compared to AO or MI alone. For lowly-coordinated AOMI (AOMI_LOCO_), where there is little overlap between the AO and MI components, the DASH proposes that visuo-motor representations for the observed and imagined actions are likely to compete as separate sensorimotor streams. This would result in similar cortico-motor activity to performing the AO or MI tasks in isolation, depending on which simulation process is prioritized based on relevance to the ongoing motor plan. For moderately-coordinated AOMI (AOMI_MOCO_), the visuo-motor representations of the observed and imagined movements may either merge or compete. This is dependent on the amount of transferable sensorimotor information between the two simulated movements and the relevance of these to the ongoing movement plan. Based on this proposition, AOMI_MOCO_ can result in more widespread pre-motor activity via merged sensorimotor streams, or similar activity to AO or MI alone through prioritization of one competing sensorimotor stream, depending on the level of coordination between the two simulated movements during this type of AOMI.

The Visual Guidance Hypothesis (VGH; Meers et al., 2020) has been proposed as an alternative account for the processes involved in *coordinative* AOMI. The VGH argues that the imagined action is prioritized, with the observed action serving as an external visual stimulus that may facilitate or disrupt the imagined action during *coordinative* AOMI. According to the VGH, the increased cortico-motor activity reported in previous literature for AOMI_HICO_ (see Chye et al., 2022 for a recent meta-analysis) is due to the observed action displaying *the same movement* and acting as a visual primer that forms a stronger motor representation for the imagined action. For AOMI_LOCO_, the VGH suggests that a motor representation will only be formed for the imagined action due to the observed action displaying *another movement* and acting as a visual distractor. Unless ignored, this will disrupt the generation and maintenance of the imagined action, meaning cortico-motor activity will be reduced compared to independent MI. In a similar vein to AOMI_LOCO_, the VGH proposes that a motor representation will only be formed for the imagined action during AOMI_MOCO_ due to the observed action displaying *another movement*. However, the observed action might act as a visual primer or visual distractor during AOMI_MOCO_ depending on the amount of transferable sensorimotor information between the two simulated movements. It is therefore possible that the observed action can both facilitate and disrupt the generation and maintenance of the imagined action, leading to either increased, similar, or decreased cortico-motor activity compared to independent MI.

Single-pulse transcranial magnetic stimulation (TMS) is the most prevalent neuroscientific modality adopted in the AOMI literature as it has high temporal resolution and can be used to determine the specific contributions for observed and imagined actions during AOMI (Chye et al., 2022). However, only three studies have used TMS to examine the neurophysiological markers for *coordinative* AOMI, and reported somewhat conflicting findings (Bruton et al., 2020; Meers et al., 2020; Grilc et al., 2024). Bruton et al. (2020) used a *coordinative* AOMI task where participants observed an index finger abduction-adduction movement while imagining the same movement with their little finger. They found that CSE was facilitated in the muscles controlling both the observed and imagined movements, when controlling for visual attention on the observed movement, supporting the DASH. Conversely, using a similar finger movement task, Meers et al. (2020) found that CSE was only facilitated for the muscles involved in the imagined finger movement, with no such facilitation for the observed finger movement during *coordinative* AOMI, in line with the VGH. In a recent TMS study, Grilc et al. (2024) utilized two different *coordinative* AOMI tasks where coordination between the observed and imagined actions was based on different movement parameters. Both *coordinative* AOMI tasks involved observation of a knee extension movement, with one task involving imagined plantarflexion of the foot, a movement that is typically coupled with a knee extension movement when the body is propelled forward or upward by the legs (i.e., coordinated based on functional coupling), and one task involving imagined dorsiflexion of the foot, an action that rotates the foot in the same direction in the sagittal plane as the rotation to the shank produced by knee extension (i.e., coordinated based on movement direction). CSE was facilitated in the muscle controlling the imagined foot movement for both *coordinative* AOMI conditions, regardless of movement parameter, but no such facilitation was identified for muscles controlling the observed knee extension, aligning with the finding of Meers et al. (2020) and the VGH. Given the contrasting support for the predictions of the DASH (Bruton et al., 2020) and VHG (Meers et al., 2020; Grilc et al., 2024), both hypotheses warrant further empirical investigation by examining the neurophysiological markers of *coordinative* AOMI.

The limited research on *coordinative* AOMI may be due, in part, to the challenge of clearly defining the parameters that govern coordination between the two simulated actions. Both Bruton et al. (2020) and Meers et al. (2020) coordinated the observed and imagined actions based on movement type (finger abduction-adduction), whereas Grilc et al. (2024) employed two AOMI conditions that coordinated the observed and imagined actions based on different movement parameters (i.e., functional coupling vs movement direction). It is important to draw from literature on AO or MI when considering possible movement parameters that might influence the ability to coordinate observed and imagined actions, and the subsequent neurophysiological markers for *coordinative* AOMI. Several studies have explored neurophysiological markers for AO and MI tasks with different force requirements, showing that CSE is increased as a function of force production during both observed (e.g., Alaerts et al., 2010, 2012; Helm et al., 2015) and imagined actions (e.g., Helm et al., 2015; Mizuguchi et al., 2013). There are mixed findings regarding movement speed during AO, with studies reporting decreased CSE (Moriuchi et al., 2014, 2017), no change in CSE (Moriuchi et al., 2017), and increased CSE (Kitamura et al., 2023) as movement speed is increased for various upper-limb movements. Given the close relationship between movement force and speed for dynamic actions, where greater force often results in greater acceleration and thus velocity, manipulating one of these parameters is of potential interest during *coordinative* AOMI.

This study aimed to test the DASH (Eaves, Riach, et al., 2016) and VGH (Meers et al., 2020) propositions for *coordinative* AOMI by comparing CSE responses during AOMI conditions where coordination between the observed and imagined actions varied based on movement speed. In this experiment, all *coordinative* AOMI conditions involved observation of a *slow-paced* single-leg sit-to-stand (SL-STS) movement but varied in the speed of the simultaneously imagined SL-STS. Specifically, AOMI_HICO_ involved simultaneous imagery of a *slow-paced* SL-STS, AOMI_MOCO_ involved simultaneous imagery of a *medium-paced* SL-STS, and AOMI_LOCO_ involved simultaneous imagery of a *fast-paced* SL-STS during observation of a *slow-paced* SL-STS, with audio sonification used to guide the imagery speed (see Castro, Osman et al., 2021; Castro, Bryjka et al., 2021).

If the propositions of the DASH (Eaves, Riach, et al., 2016) are accurate, it is hypothesized that: H1; CSE facilitation will be increased for the AOMI_HICO_ condition at T3 (i.e., the stimulation timepoint corresponding to peak EMG activity for the *slow-paced* SL-STS) due to the merging of sensorimotor streams for AO and MI of the same action and movement speed. H2; CSE facilitation will be increased for the AOMI_MOCO_ condition at both T3 and T2 (i.e., the stimulation timepoint corresponding to peak EMG activity for the *medium-paced* SL-STS) due to the maintenance of sensorimotor streams for AO and MI of the same action with similar movement speeds. H3; CSE facilitation will not be increased for the AOMI_LOCO_ condition at any stimulation timepoint due to the competition between sensorimotor streams for AO and MI of the same action with different movement speeds.

If the propositions of the VGH (Meers et al., 2020) are accurate, it is hypothesized that: H1; CSE facilitation will be increased for the AOMI_HICO_ condition at T3 because the observed action will enhance the motor representation formed for the imagined action when the same movement type and speed are employed for both AO and MI simultaneously. H2; CSE facilitation will be increased for the AOMI_MOCO_ condition at T2 due to prioritization of the imagined action during simultaneous AO and MI of the same action with similar movement speeds. H3; CSE facilitation will be increased for the AOMI_LOCO_ condition at T1 due to prioritization of the imagined action during simultaneous AO and MI of the same action with different movement speeds.

## Methods

### Participants

Based on previous AOMI studies employing TMS (e.g., Bruton et al., 2020; Grilc et al., 2024; Wright et al., 2018), twenty-one healthy adults aged 21-43 years (14 male, M_age_ = 29.29 ± 6.78 years) with normal or corrected-to-normal vision took part in this study. All participants underwent screening and provided written informed consent prior to data collection. The screening included completion of the TMS Adult Safety Screen (TASS; Keel et al., 2001) and the Movement Imagery Questionnaire - 3 (MIQ-3; Williams et al., 2012). All participants were safe to take part in the study based on responses to the TASS. MIQ-3 scores indicated that the final study sample could easily generate internal visual (6.14 ± 0.66), external visual (5.71 ± 1.27), and kinesthetic (5.57 ± 1.22) imagery, confirming their ability to complete the experimental tasks.

### Experimental Design

This study was conducted in accordance with the ethical guidelines and approval of the ethical committee at the host university (ethical approval number LSC 21-346). All study procedures are reported using parts A, B and C of the Guidelines for Reporting Action Simulation Studies checklist (Moreno-Verdú et al., 2024) and experimental code and materials can be accessed via the Open Science Framework: https://osf.io/5vjwe/. A repeated measures design was employed where participants completed six conditions, split into three control conditions and three experimental conditions, with all conditions repeated across two testing sessions. The three control conditions were: (i) a baseline (BL) condition where participants looked at a fixation cross placed on a static image depicting the SL-STS from the video stimuli (50% of trials) or a blank background (50% of trials), (ii) an action observation (AO) condition where participants were required to observe a video of a model performing a *slow-paced* SL-STS movement, and (iii) a motor imagery (MI) condition where participants were required to imagine the feelings and sensations associated with performing a *slow-paced* SL-STS movement. The three experimental conditions were: (iv) a highly-coordinated AOMI (AOMI_HICO_) condition where participants were asked to simultaneously observe a *slow-paced* SL-STS movement while imagining the feelings and sensations of performing the same *slow-paced* SL-STS movement, (v) a moderately-coordinated AOMI (AOMI_MOCO_) condition where participants were asked to simultaneously observe a *slow-paced* SL-STS movement while imagining the feelings and sensations of performing a *medium-paced* SL-STS movement, and (vi) a lowly-coordinated AOMI (AOMI_LOCO_) condition where participants were asked to simultaneously observe a *slow-paced* SL-STS movement while imagining the feelings and sensations of performing a *fast-paced* SL-STS movement.

The six conditions were presented in a semi-randomized order across participants (see Figure 1 for a visual depiction of the experimental design), consistent with previous TMS studies investigating different forms of AOMI (e.g., Bruton et al., 2020; Grilc et al., 2024). The BL condition was delivered across five blocks of six trials (i.e., blocks 1, 3, 5, 7, 9) to account for a possible ordering effect across the experiment by recording comparator MEP amplitude data before each condition containing movement simulation tasks. The AO condition was presented as block 2, before any MI instructions were provided, to reduce the likelihood of participants engaging in spontaneous or deliberate MI during this condition (see Bruton et al., 2020). The MI condition was presented as block 4, after AO, as it was deemed necessary to have first exposed participants to visual stimuli of a SL-STS to facilitate their MI of the action due to their naivety with this movement. The three experimental conditions involving *coordinative* AOMI tasks were presented last (i.e., blocks 6, 8, and 10), with the order of these counterbalanced across the study sample to avoid the possibility of an ordering effect for these conditions.

**Figure 1.**
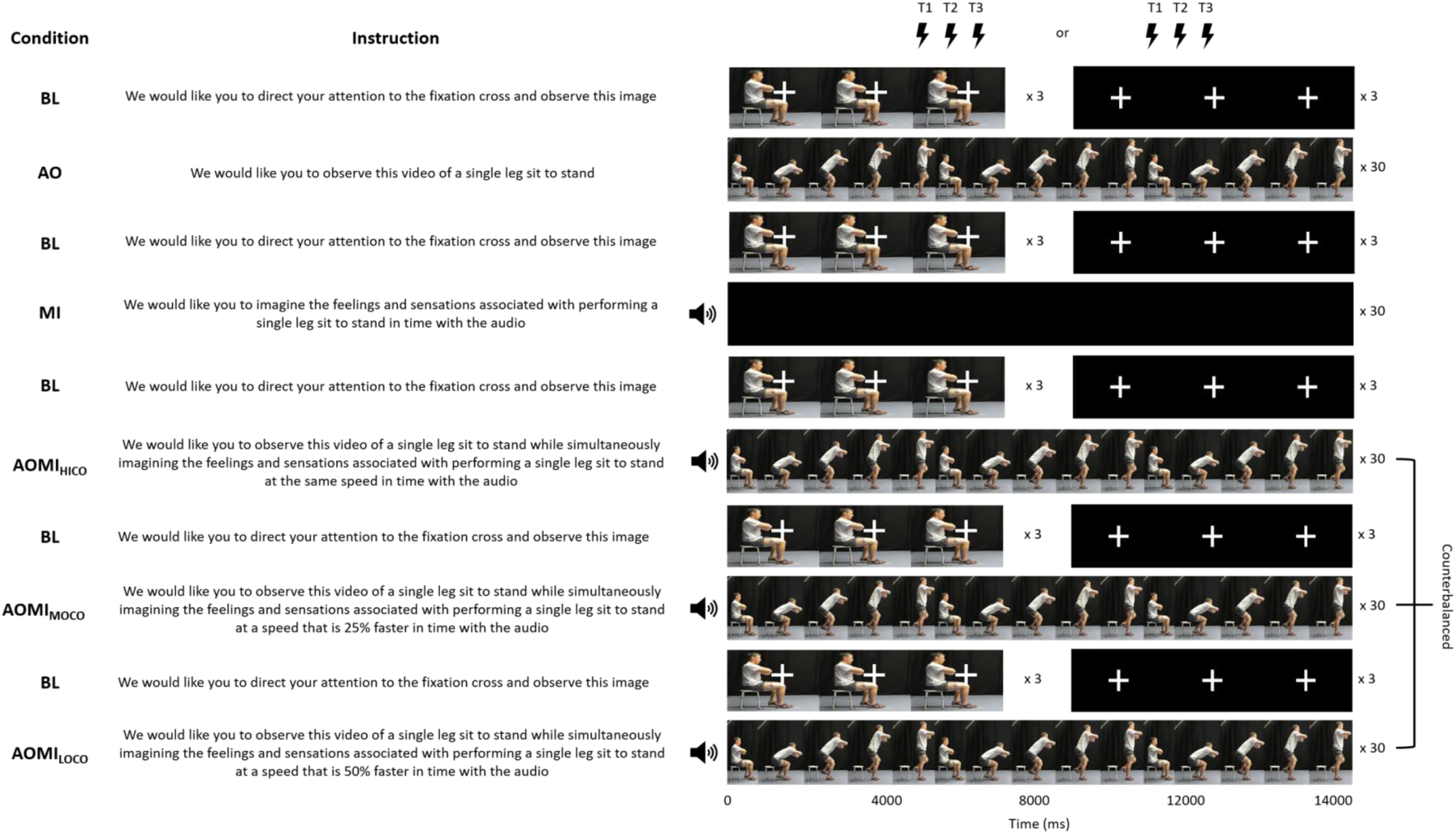
A Visual Representation of the Ten Experimental Blocks Completed During Each Testing Session. *Note:* The TMS was delivered at the point of peak EMG activity in the knee extensor muscle group for the models performing the *fast-paced* (T1), *medium-paced* (T2), and *slow-paced* (T3) SL-STS for either the second or third cycle of every AOMI trial, and at the same time point for trials in control conditions. The ordering of the TMS delivery was randomized and counterbalanced across trials for each experimental block. The model depicted in this figure is an author of this study.

### Target Movement

Rising from a chair, often referred to as sit-to-stand (STS), is a task that requires strength, balance, and coordination. In elderly individuals, especially those with functional limitations, STS performance is associated with various measures of physiological function (Lord et al., 2002; Schenkman et al., 1996) and the demands of this movement can approach or exceed strength capacity (Hughes et al., 1996). This poses a high risk of falls (Robinovitch et al 2013), making it a task of significant importance, especially in the maintenance of independent living for older adult populations (Slaughter et al., 2015). Although a normal STS is a relatively unchallenging task for young individuals, it becomes challenging in a manner that resembles the differences between young and older adults performing a normal STS if performed on a single leg (SL-STS). For example, young adults exhibit a significantly longer movement duration during SL-STS relative to normal STS, with movement duration becoming negatively rather than positively correlated with strength for this adapted movement (Thongchoomsin et al 2020). For these reasons, the SL-STS was chosen as the target movement in this study.

### Stimuli Development

Timings for the *slow-, medium-* and *fast-paced* SL-STS movements depicted in the video and audio stimuli used in this study were determined through pilot testing given the novelty of the selected target movement. Five pilot participants (4 male, 1 female) completed five successful trials of the SL-STS under instructions to move (i) as slowly as possible (i.e., *slow-paced*), (ii) at a ‘normal’ pace (i.e., *medium-paced*), and (iii) as quickly as possible (i.e., *fast-paced*). During these trials, a marker on the C7 vertebra was tracked to identify movement start and end points using a 12-camera Vicon Vantage motion capture system (Vicon, Oxford, UK) sampling at 100Hz. Surface electromyography (EMG) activity was recorded for the vastus lateralis (VL), vastus medialis (VM) and rectus femoris (RF) quadricep muscles using a single bipolar silver-chloride gel-electrode configuration on each muscle (2-cm diameter, 2-cm inter-electrode distance; Dual Electrode, Noraxon, Arizona, U.S.A.) to detect the point of peak EMG activity for the *slow*-, *medium-* and *fast-paced* SL-STS movements. Quadricep muscles were selected as the target muscles because they are a large contributor to forces during the STS (Ahmadi-Ahangar et al., 2018; Miyoshi et al., 2005). On average across the pilot participants, the *slow-paced* SL-STS trial lasted 3.76 ± 1.23-s, the *medium-paced* SL-STS trial lasted 2.34 ± 0.38-s, and the *fast-paced* SL-STS trial lasted 1.69 ± 0.29-s. Since the average *medium-paced* SL-STS trial had a duration which was closer to the duration of the *fast-paced* SL-STS trial, it was decided that the *medium-paced* SL-STS would be best represented by the midpoint between the *slow* and *fast-paced* SL-STS (i.e., 2.73-s).

Once the timings for the three SL-STS speeds were established, one male and one female model were recruited for development of the video stimuli. The models were video recorded performing the *slow-, medium-* and *fast-paced* SL-STS movement in the sagittal plane (see Figure 2) using a Panasonic camera (HC-W570, 1920×1080 resolution and 50 frames per second). A metronome was used so that both models could match the selected timings for all three movement speeds as closely as possible.

**Figure 2.**
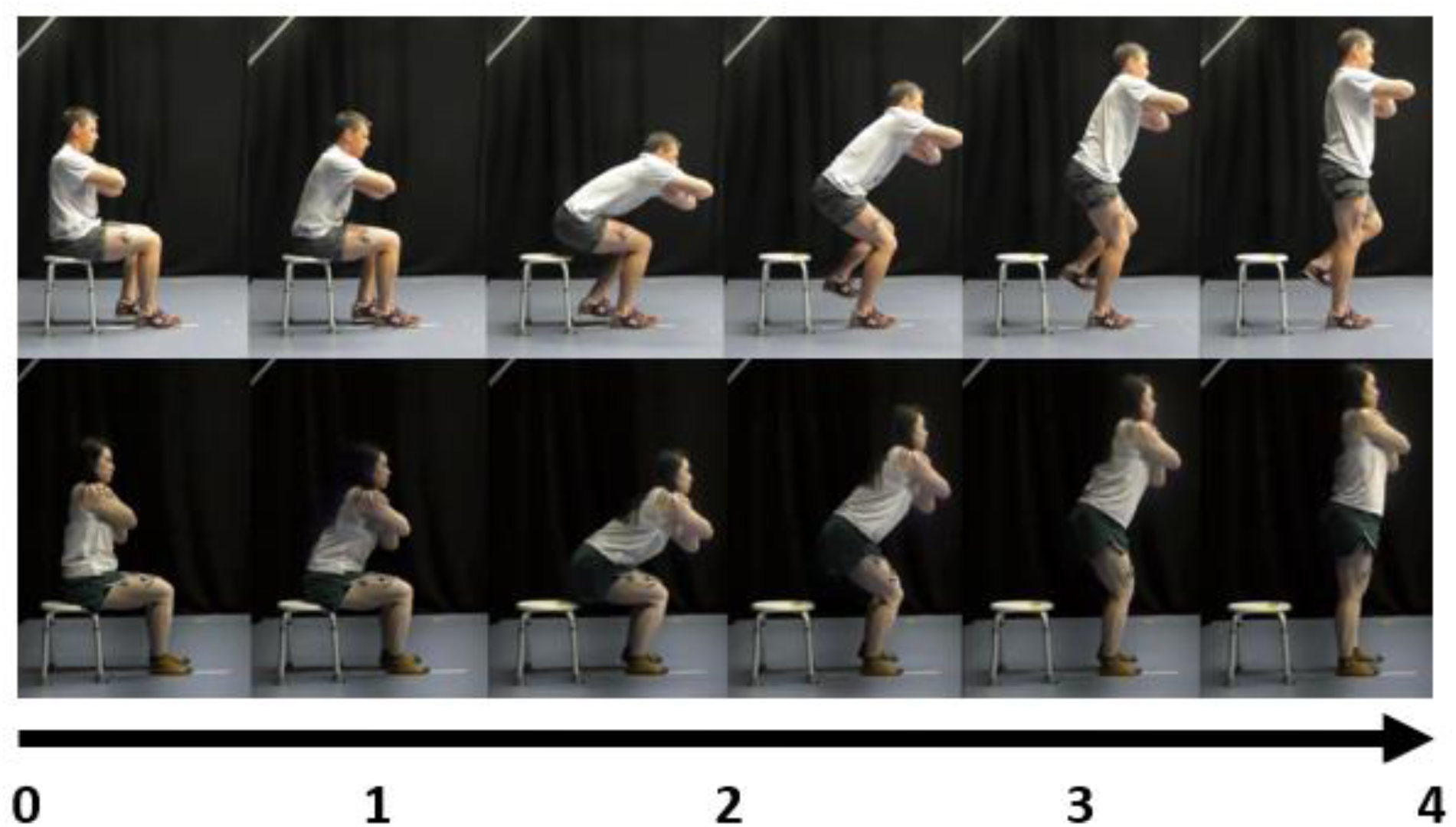
A visual depiction of the male (top) and female (bottom) models performing the *slow-paced* SL-STS used as the video stimuli in the AO control condition and AOMI experimental conditions used in this study. *Note:* The timing differed across the models, with the male model taking 3740ms and the female model taking 3660ms to complete the *slow-paced* SL-STS used in the video stimuli. The models depicted in this figure are both authors of this study.

After reviewing footage captured for the models, the research team agreed upon each model’s best trial for the different SL-STS speeds by considering the time-discrepancy between the target and executed trial durations, and the perceived smoothness of the executed SL-STS movement. The stimulation timings were calculated based on the EMG data collected from the best trials of the two models. Using a bespoke MATLAB (R2020a, The MathWorks) script, the peak EMG activity for each muscle at the three different speeds was identified for both models. The absolute peak time was then determined and normalized as a percentage of the total movement duration. These normalized values were averaged across muscles for each speed and model. Movement durations for slow, medium, and fast speeds were defined based on the slow movement duration, with the medium duration calculated as 75% of the slow movement time and the fast duration defined as 50% of the slow movement time respectively for each model. Finally, the stimulation times were derived by relating the average normalized peak activity to these predefined durations (see https://osf.io/n7p2a for formula used). The calculated movement duration times were used to produce movement sonification files on the open-source software Audacity (Audacity Team version 2.4.1) to guide MI timing during conditions where this was required (i.e., the MI control condition and three experimental conditions) in-line with the procedures of Castro and colleagues (Castro, Bryjka, et al., 2021; Castro, Osman, et al., 2021).

## Experimental Procedure

### Surface Electromyography Preparation and Recording

Prior to EMG placement, the skin was prepared by shaving and cleaning the area where EMG electrodes were placed. The EMG system adopted for stimuli development was used to measure EMG activity for three KE muscles of the participants’ right leg during the main experimental protocol. The EMG sensors (DTS-EMG sensors, Noraxon, USA) and electrodes were attached to the muscle belly of the target muscles based on the SENIAM (Hermens et al., 1999) guidelines. Noraxon wireless sensors (bandwidth of 20–450 kHz, 92 dB common mode rejection ratio and > 1015 Ω input impedance) transmitted the EMG signals to a desktop receiver (TeleMYO DTS EMG; Noraxon, Arizona, USA), and these were sampled at 2 kHz via an analogue-to-digital convertor (Micro 1401–3) and desktop PC utilizing Spike2 software (Cambridge Electronic Design, Cambridge, UK).

### Transcranial Magnetic Stimulation Preparation

#### Optimal Scalp Position

The TMS preparation procedures adhered to methodological guidelines, as recorded via a methodological reporting checklist for TMS experiments (Chipchase et al., 2012) and the methods employed in this study were adapted from established protocols (Grilc et al., 2024).

Participants were asked to wear a tight-fitting polyester cap with the center of the scalp (i.e., Cz), defined by the intersection of lines from inion to nasion and from left to right tragus, measured and marked as the top-right most corner of a 3×3cm grid drawn for identification of the optimal scalp position (OSP). A single-pulse double-cone coil TMS (110 mm diameter) connected to a Magstim 200^2^ monophasic magnetic stimulator was originally placed at Cz and oriented to direct current flow from anterior to posterior. The TMS coil was manually moved around the 3×3cm grid in 0.5cm intervals posterior and lateral from Cz, and four stimulations were delivered at each site to identify the OSP that produced the largest and most consistent MEP amplitudes across the three KE muscles. OSP sites ranged from 0.5-2.5cm posterior and 0-1.5cm lateral to Cz, with the median OSP coordinates recorded as 1cm posterior and 0.5cm lateral across the study sample.

#### Resting Motor Threshold

The resting motor threshold (RMT; excitability of the KE muscle representation of the motor cortex at rest) was determined following the guidelines of Rossini et al. (2015). This involved gradually reducing the stimulation intensity in 5% increments from the participants’ OSP intensity until five out of ten trials produced MEP amplitudes exceeding 50 μV in at least two out of three KE muscles. Consistent with previous TMS literature on AOMI, the experimental stimulation intensity was set at 110% of the RMT (Bruton et al., 2020; Grilc et al., 2024; Wright et al., 2018) to minimize direct wave stimulation (Loporto et al., 2013). The mean RMT was 64 ± 11% and the mean experimental stimulation intensity was 71 ± 12% of the maximum stimulator output, both similar values to that of a recent TMS experiment investigating AOMI in lower-leg muscles (see Grilc et al., 2024).

#### Experimental Setup

The participants were seated comfortably on an isokinetic dynamometer chair (HUMAC NORM, CSMi, Stoughton, MA) in a dimly-lit Biomechanics Laboratory at the host university, with their upper body loosely strapped in place to reduce movement or repositions during the experimental block and maintain a consistent viewing position between blocks. The video stimuli were presented on an 80” adjustable screen (LED Interactive Multi-Touch Display, model 86GT-4K, SHENZHEN Hitevision Technology Co., Ltd.) placed 130cm directly in front of the participant (Figure 3) using DMASTR DMDX display software (Forster & Forster, 2003).

**Figure 3.**
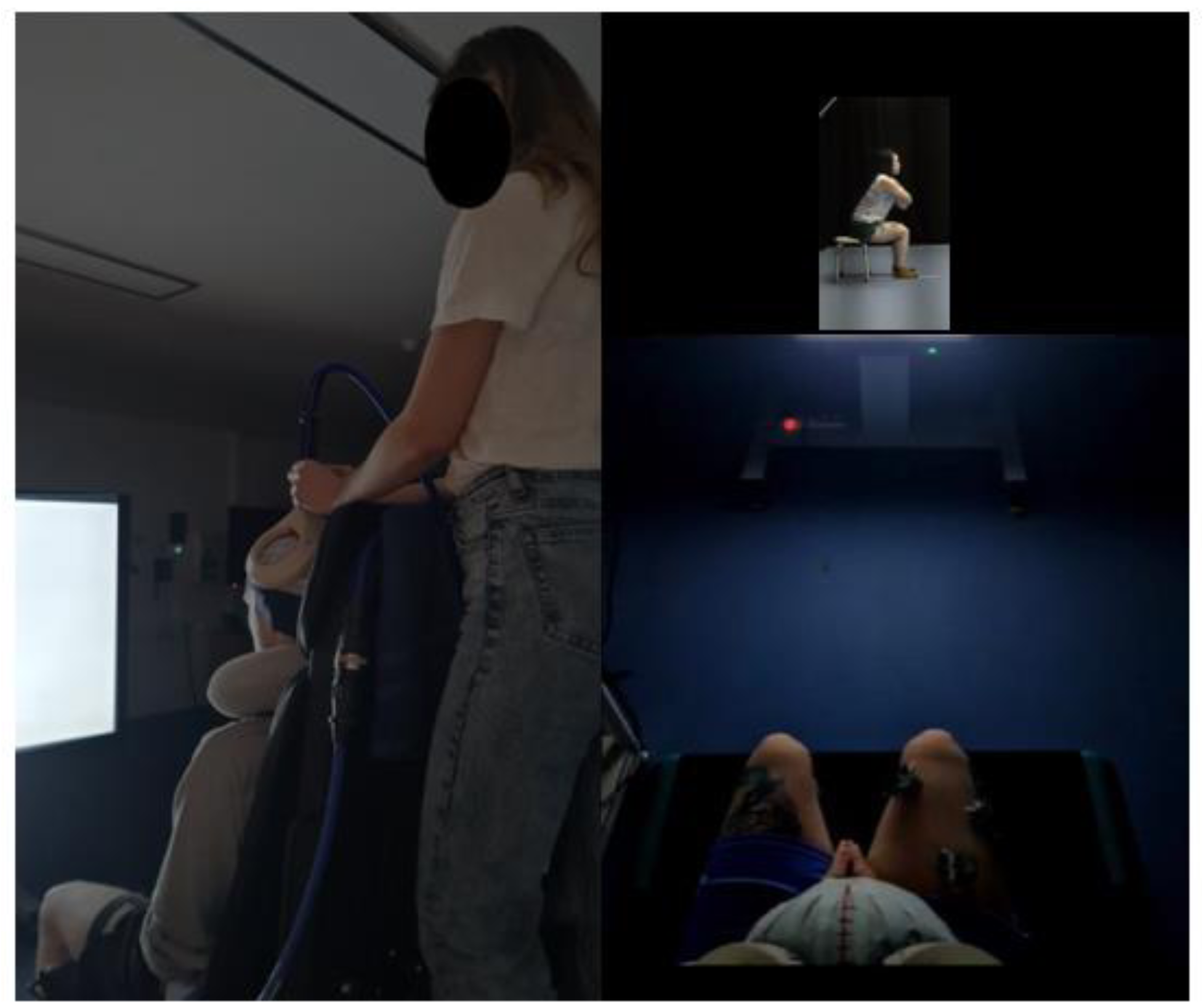
Experimental setup from two perspectives. The left image shows the experimenter positioning the TMS coil over the participant seated on the Cybex dynamometer, facing the screen. The right image offers an overhead view, approximating the participant’s perspective with the coil in place. The people depicted in this figure are all authors of this study.

#### Familiarization

A familiarization phase was incorporated for all conditions where MI was instructed (i.e., MI, AOMI_HICO_, AOMI_MOCO_, AOMI_LOCO_) to aid the participants’ engagement with the upcoming experimental tasks. During this phase, participants watched five repetitions of a model performing the SL-STS at the required speed whilst seated on an adjustable stool with arms placed diagonally across their chest. After this, the participants physically practiced the movement for five repetitions, matching their speed to the speed of the model in the demonstration video. A further five physical repetitions were then completed, with audio sonification added to the video of the model performing the SL-STS at the speed to be imagined. The final step involved the participants receiving instructions to engage with MI of the SL-STS in time with the audio sonification of the movement whilst being presented with a video of a *slow-paced* SL-STS for the experimental conditions, or image of a blank screen for the MI condition.

#### Experimental Protocol

The experimental protocol was repeated across two 3-hour testing sessions, at least 48 hours apart. This design ensured the collection of sufficient data points for the study while minimizing participant fatigue and discomfort. In each testing session, the participants completed ten experimental blocks consecutively, with the five BL blocks lasting 90-s and the five main experimental blocks (i.e., AO, MI, AOMI_HICO_, AOMI_MOCO_, AOMI_LOCO_) lasting 450-s per block. A 180-srest period was included after each of the main experimental blocks, and participants were encouraged to leave the testing chair to prevent eye strain and muscular discomfort before familiarizing themselves with the stimuli and task for the next experimental block. Prior to beginning the experiment, participants were asked to read the on-screen instructions carefully, refrain from voluntary movement during the experimental blocks, and to attend fully to the stimuli presented. Understanding of these instructions was verbally confirmed before the start of each block. The 30-trial main experimental block incorporated a rest period after each set of 10 trials. Written and verbal reminders of the specific instructions for the condition were provided before the participant completed each set of 10 trials. All trials were displayed on the LCD screen using DMASTR DMDX display software (Forster & Forster, 2003) and lasted 10980ms for female participants and 11220ms for male participants. For the AO control condition and three AOMI experimental conditions (see Figures 1 and 2 for stimuli and instructions), the trials showed three repetitions of a *slow-paced* SL-STS (3660ms per cycle for females, 3740ms per cycle for males). When observing the *slow-paced* SL-STS, participants were instructed to simultaneously imagine the feelings and sensations associated with a slow-(AOMI_HICO_), *medium-* (AOMI_MOCO_), or *fast-paced* SL-STS (AOMI_LOCO_) using audio sonification files lasting 100%, 75% and 50% of the *slow-paced* SL-STS duration for respective guidance. For the MI condition, the participants observed a blank screen and engaged with MI following the same instructions and audio sonification as the AOMI_HICO_ condition.

#### TMS Data Collection

Using a script run through Spike 2 software, a single TMS pulse was delivered once per trial during the second (female = 3661-7320ms, male = 3741-7580ms) or third (female = 7321-10980ms, male = 7581-11220ms) cycles of the observed *slow-paced* SL-STS movement used in the three experimental and AO conditions, and at the corresponding time points for the trials for the BL and MI control conditions. Participants were stimulated during two cycles of the SL-STS movement to reduce the predictability of the stimulation and subsequent anticipatory behavior of the participants (Loporto et al., 2012). TMS stimulation timepoints represented the point of peak EMG activity for the two models when physically performing the different paced SL-STS trials used to produce the video and audio stimuli for this study. TMS stimulation timepoints corresponding to the *fast-* (T1; female = 450ms, male = 550ms after cycle onset), *medium-* (T2; female = 610ms, male = 720ms after cycle onset) and *slow-paced* SL-STS (T3; female = 800ms, male = 1300ms after cycle onset) were randomly ordered and counterbalanced within-conditions. A 3-s transition period was implemented between experimental trials to maintain an inter-stimulus interval greater than 10-s to let the effects of the preceding TMS pulse diminish (Chen et al., 1997).

#### Social Validation Data Collection

Ratings of perceived MI ability were recorded using a 7-point Likert scale from 1 (*very easy to feel*) to 7 (*very hard to fee*l) at the end of the familiarization phase, during the break periods between each block of 10 trials, and at the end of the experiment for the conditions involving engagement with MI processes (i.e., MI, AOMI_HICO_, AOMI_MOCO_, AOMI_LOCO_). After finishing the TMS data collection for the second testing session, participants recorded social validation data in a similar fashion to recent AOMI experiments using TMS (e.g., Bruton et al., 2020; Grilc et al., 2024; Riach et al., 2018). Initially, participants recorded responses on a bespoke social validation questionnaire, where they rated perceived sex and ownership of the modeled SL-STS movements displayed in the conditions with an AO component (i.e., AO, AOMI_HICO_, AOMI_MOCO_, AOMI_LOCO_) using three 5-point Likert scales from 1 (*totally disagree*) to 5 (*totally agree*). Participants then took part in a semi-structured social validation interview designed to understand their experiences across the different conditions and check for compliance with the intended manipulations.

## Data Analysis

### TMS Data Processing

Peak-to-peak MEP amplitudes were recorded from three KE muscles (VM, VL and RF) of the participants’ right leg on a trial-by-trial basis and averaged across all successful trials for T1, T2 and T3 TMS stimulation timepoints per condition from both testing sessions. After extracting the data from Spike 2, and separating the data based on TMS stimulation timepoint, a two-part screening process was used to determine successful trials. First, trials were screened for the presence of an MEP response, and any trials that did not evoke an MEP amplitude of sufficient magnitude (i.e., 50 μV) were removed. It is well-established that MEP amplitudes are increased for a target muscle if the EMG activity in that muscle is above resting state levels at, or immediately prior to, the time of TMS stimulation (Devanne et al., 1997; Hess et al., 1987). To control for this, EMG activity was recorded for 200 ms prior to the delivery of each TMS pulse and any trials where the EMG amplitude exceeded normal baseline values (mean ± 2.5 SD) for that TMS stimulation timepoint and condition were removed (e.g., Riach et al., 2018; Wright et al., 2014; 2018). For the study sample, a mean value of 4.71 (± 3.93) trials for the VM muscle, 6.66 (± 6.05) trials for the VL muscle, and 2.35 (± 2.59) trials for the RF muscle were removed per TMS stimulation timepoint and condition. This resulted in the sample having a mean value of 19.22 (± 4.69) trials with an uncontaminated MEP response in one or more KE muscles, providing a reliable estimate of CSE per TMS stimulation timepoint and condition (Cuypers et al., 2014). On a muscle-by-muscle basis, the raw MEP amplitude data for successful trials was normalized using a *z*-score transformation to account for the large intra- and inter-participant variability in MEP amplitudes at rest (e.g., Bruton et al., 2020; Grilc et al., 2024; Wright et al., 2014). For each participant, this procedure involved standardizing the MEP amplitude value recorded for each successful trial against all other MEP amplitude values recorded for successful trials for that TMS stimulation timepoint across the two testing sessions. The mean amplitude for all successful trials at each TMS stimulation timepoint (i.e., T1, T2, T3) was represented by a value of zero, and values for each condition were denoted by how many standard deviations that condition was above or below the mean of all conditions. Once the *z*-score transformation was complete, the mean *z*-score MEP amplitude value was recorded for the three KE muscles on a trial-by-trial basis. This mean was calculated from three KE muscles for 70.41%, two KE muscles for 19.83%, and one KE muscle for 9.76% of successful trials. For all participants, a mean *z*-score MEP amplitude value was calculated per condition at each TMS stimulation timepoint for analysis.

### TMS Data Analysis

The *z*-score MEP amplitude data for the KE muscles was normally distributed at each TMS stimulation timepoint, permitting the use of analysis of variance (ANOVA) statistical tests. Three separate one-way repeated measure ANOVA tests with six levels (Condition: BL, AO, MI; AOMI_HICO_; AOMI_MOCO_; AOMI_LOCO_) were run using the rstatix package (Kassambara, 2023) and the data was visualized using the ggplot2 package (Wickham, 2016) in R studio statistical software (version 4.3.2). Bonferroni contrasts were used for post-hoc pairwise comparisons. Outlier analysis was conducted using interquartile range values, and data points identified as outliers were removed from the three ANOVA analyses (T1 = 2, T2 = 2, T3 = 3 outliers). To test for a potential ordering effect across the experiment, a one-way repeated measure ANOVA test with five levels (BL block: block 1, block 3, block 5, block 7, block 9) was conducted for *z*-score MEP amplitude data recorded in BL experimental blocks. For this comparison, the *z*-score transformation was conducted based on the data from the 60 BL trials collected across the two testing sessions. A one-way repeated measures ANOVA analysis performed on the BL condition *z*-score MEP amplitude data revealed no significant main effect of block, F_(4, 72)_ = 1.01, *p* = .41, η_p_^2^ = .05.

### Social Validation Data Analysis

A one-way repeated measures ANOVA with four levels (Condition: MI; AOMI_HICO_; AOMI_MOCO_; AOMI_LOCO_) was run on the perceived MI ability data collected from the social validation questionnaire to investigate differences across the conditions with MI processes. An independent samples t-test (Participant Sex: male; female) was run on the perceived body ownership, masculinity, and femininity data collected from the social validation questionnaire to investigate differences in male and female participants’ ratings of the movements displayed across the conditions with AO processes. Interview data was transcribed, coded, and grouped on a question-by-question basis to offer detailed explanations for the MI ability ratings from the social validation questionnaire.

## Results

### MEP Amplitude Data

One-way repeated measures ANOVA analyses performed on the Z-score MEP amplitude data revealed a significant large main effect of condition at T1, F_(5, 90)_ = 8.28, *p* < .001, η_p_^2^ = .31, T2, F_(3.04, 54.69)_ = 8.32, *p* < .001, η_p_^2^ = .31, and T3, F_(5, 85)_ = 18.4, *p* < .001, η_p_^2^ = .51, TMS stimulation timepoints. Post-hoc pairwise comparisons between the experimental conditions (i.e., AOMI_HICO_, AOMI_MOCO_, AOMI_LOCO_) and control conditions (i.e., BL, AO, MI) across the three TMS stimulation timepoints are reported below.

### Highly-Coordinated AOMI (AOMI_HICO_)

Post-hoc pairwise comparisons at T3 (Figure 4), the TMS stimulation timepoint corresponding to the *slow-paced* SL-STS, showed that Z-score MEP amplitudes were significantly larger in the AOMI_HICO_ condition compared to the BL (*p* < .001) and AO (*p* < .001) conditions, but not the MI condition (*p* = .29).

**Figure 4.**
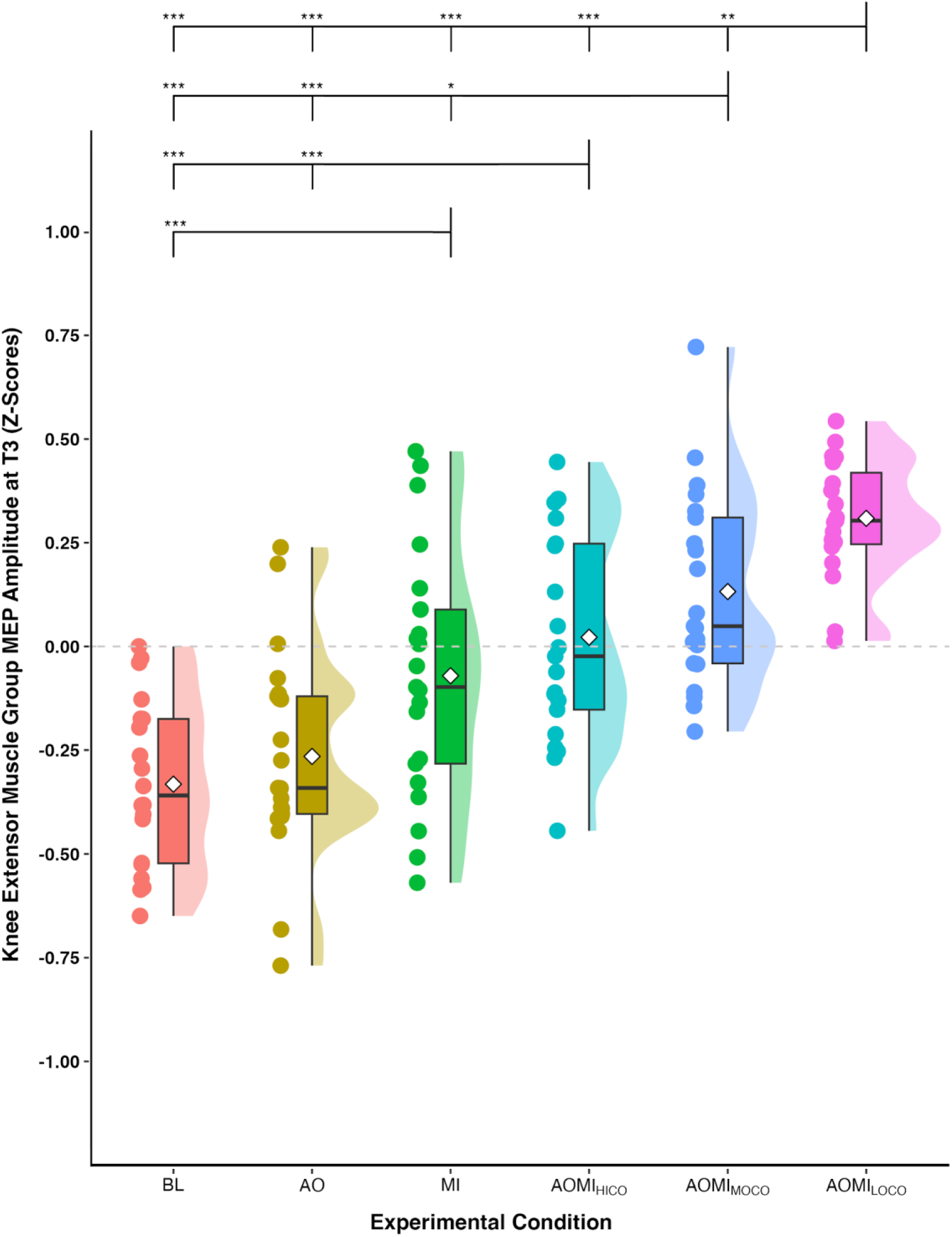
Box and violin plot with raw data points displaying z-score normalized MEP amplitudes from the knee extensor muscle group at the stimulation timepoint corresponding to the *slow-paced* SL-STS (T3) for the three control and three experimental conditions. *Note:* Thick horizontal black lines represent the median average and white diamonds represent the mean average for each box plot. Individual participant data points are represented by circular markers. BL – baseline; AO – action observation; MI – motor imagery; AOMI_HICO_ – highly-coordinated action observation and motor imagery; AOMI_MOCO_ – moderately-coordinated action observation and motor imagery; AOMI_LOCO_ – lowly-coordinated action observation and motor imagery; \**p* < .05, \*\**p* < .01, \*\*\**p* < .001.

Post-hoc pairwise comparisons at T2 (Figure 5), the TMS stimulation timepoint corresponding to the *medium-paced* SL-STS, showed no significant differences between the AOMI_HICO_ condition and the three control conditions (*p*s > .05). Post-hoc pairwise comparisons at T1 (Figure 6), the TMS stimulation timepoint corresponding to the *fast-paced* SL-STS, showed that Z-score MEP amplitudes were significantly larger in the AOMI_HICO_ condition compared to the BL (*p* = .01) and AO (*p* < .001) conditions, but not the MI condition (*p* = .34).

**Figure 5.**
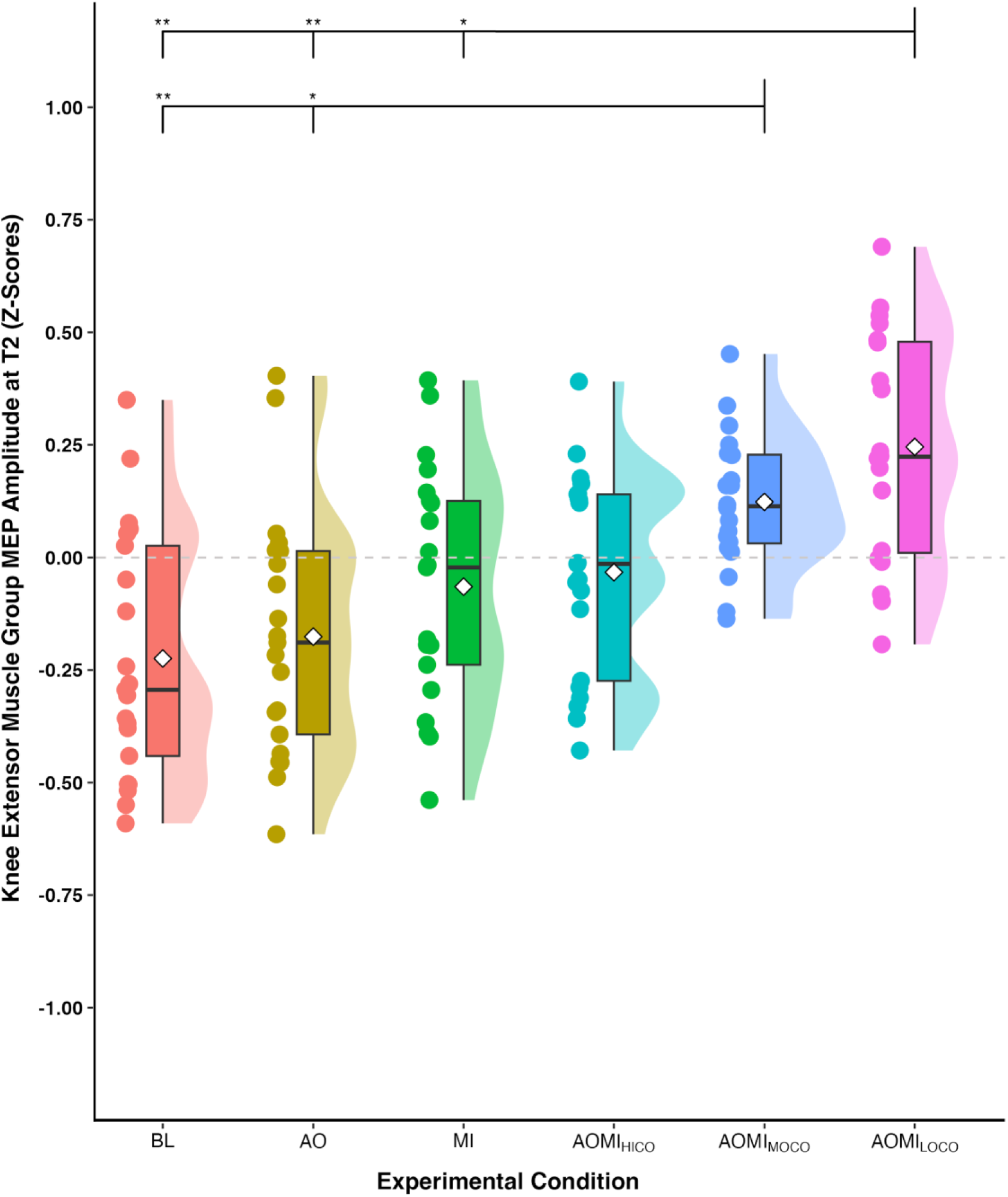
Box and violin plot with raw data points displaying z-score normalized MEP amplitudes from the knee extensor muscle group at the stimulation timepoint corresponding to the *medium-paced* SL-STS (T2) for the three control and three experimental conditions. *Note:* Thick horizontal black lines represent the median average and white diamonds represent the mean average for each box plot. Individual participant data points are represented by circular markers. BL – baseline; AO – action observation; MI – motor imagery; AOMI_HICO_ – highly-coordinated action observation and motor imagery; AOMI_MOCO_ – moderately-coordinated action observation and motor imagery; AOMI_LOCO_ – lowly-coordinated action observation and motor imagery; \**p* < .05, \*\**p* < .01.

**Figure 6.**
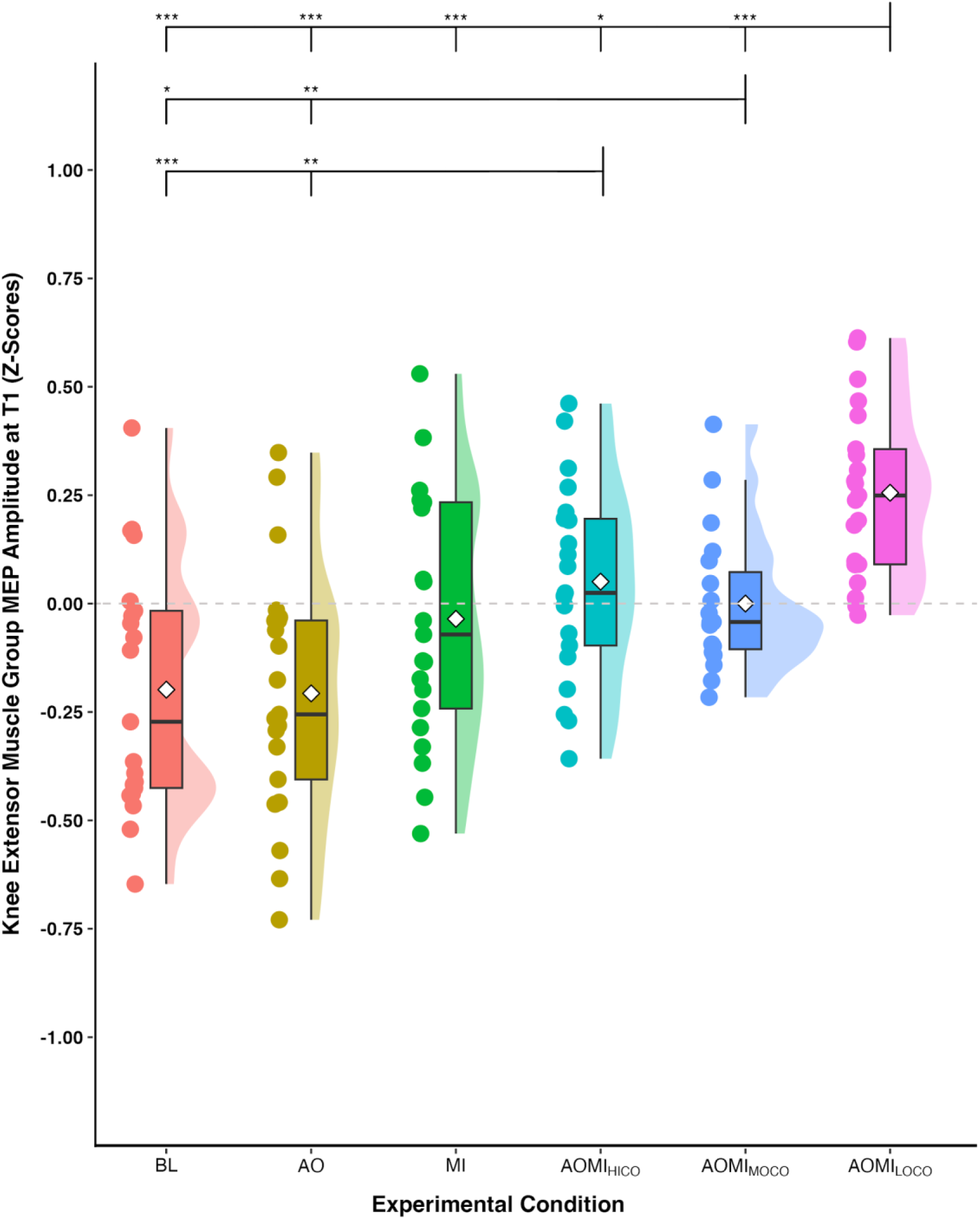
Box and violin plot with raw data points displaying z-score normalized MEP amplitudes from the knee extensor muscle group at the stimulation timepoint corresponding to the *fast-paced* SL-STS (T1) for the three control and three experimental conditions. *Note:* Thick horizontal black lines represent the median average and white diamonds represent the mean average for each box plot. Individual participant data points are represented by circular markers. BL – baseline; AO – action observation; MI – motor imagery; AOMI_HICO_ – highly-coordinated action observation and motor imagery; AOMI_MOCO_ – moderately-coordinated action observation and motor imagery; AOMI_LOCO_ – lowly-coordinated action observation and motor imagery; \**p* < .05, \*\**p* < .01, \*\*\**p* < .001.

### Moderately-Coordinated AOMI (AOMI_MOCO_)

Post-hoc pairwise comparisons at T2 (Figure 5), the TMS stimulation timepoint corresponding to the *medium-paced* SL-STS, showed that Z-score MEP amplitudes were significantly larger in the AOMI_MOCO_ condition compared to the BL (*p* = .01) and AO (*p* = .02) conditions, but not the MI condition (*p* = .07). Post-hoc pairwise comparisons at T3 (Figure 4), the TMS stimulation timepoint corresponding to the *slow-paced* SL-STS, showed that Z-score MEP amplitudes were significantly larger in the AOMI_MOCO_ condition compared to the BL (*p* < .001), AO (*p* < .001) and MI (*p* = .02) conditions. Post-hoc pairwise comparisons at T1 (Figure 6), the TMS stimulation timepoint corresponding to the *fast-paced* SL-STS, showed that Z-score MEP amplitudes were significantly larger in the AOMI_MOCO_ condition compared to the BL (*p* = .04) and AO (*p* = .01) conditions, but not the MI condition (*p* = .56).

### Lowly Coordinated-AOMI (AOMI_LOCO_)

Post-hoc pairwise comparisons at T1 (Figure 6), the TMS stimulation timepoint corresponding to the *fast-paced* SL-STS, showed that Z-score MEP amplitudes were significantly larger in the AOMI_LOCO_ condition compared to the BL (*p* < .001), AO (*p* < .001), and MI conditions (*p* < .001). Post-hoc pairwise comparisons at T2 (Figure 5), the TMS stimulation timepoint corresponding to the *medium-paced* SL-STS, showed that Z-score MEP amplitudes were significantly larger in the AOMI_LOCO_ condition compared to the BL (*p* = .01), AO (*p* = .01), and MI conditions (*p* = .02). Post-hoc pairwise comparisons at T1 (Figure 6), the TMS stimulation timepoint corresponding to the *slow-paced* SL-STS, showed that Z-score MEP amplitudes were significantly larger in the AOMI_LOCO_ condition compared to the BL (*p* < .001), AO (*p* < .001), and MI conditions (*p* < .001).

### Social Validation Data

#### Questionnaire Responses

A one-way repeated measures ANOVA analysis performed on the perceived MI ability data revealed a significant effect of condition F_(3, 60)_ = 14.23, *p* < .001, η_p_^2^ = .14. Post-hoc pairwise comparisons showed that perceived MI ability scores were significantly larger in the AOMI_HICO_ condition compared to the MI (*p* < .001), AOMI_MOCO_ (*p* < .001), and AOMI_LOCO_ (*p* < .001) conditions. All participants felt like they were looking at a same-sex model, with male participants recording a mean response of 1.67 (±0.50) indicating total agreement that the model they watched was male, and female participants recording a mean response of 1.83 (±0.41) indicating total agreement that the model they watched was female. Despite the acknowledgement that the models were the same-sex as the participants, only six participants (28.57%) felt like they were watching their own performance of the *slow-paced* SL-STS.

#### Interview Responses

The interview data provided additional detail about the preferences and strategies adopted by the participants across the experimental conditions. Participants provided mixed responses about perceived MI ability across the conditions where MI was instructed. Sixteen participants (76.19%) suggested MI was easiest in the AOMI_HICO_ condition, with most participants outlining that the synchrony between the observed video and the audio sonification of the SL-STS reduced the difficulty of the cognitive task and facilitated their imagery of the movement (e.g., “It was playing at the correct speed, so I didn’t even need to do that [synchronize], so all my attention could be focused on imagining the feeling and sensations” [participant 3]). Conversely, thirteen participants (61.90%) indicated that MI was most difficult in the AOMI_MOCO_ condition and eight participants (38.10%) reported that AOMI_LOCO_ was the most challenging condition in terms of MI ability. Participants who found MI most difficult during AOMI_MOCO_ noted that the close overlap between the speed of the AO and MI processes made it harder to time and maintain the imagined *medium-paced* SL-STS (e.g., “I think the one where I imagined the middle speed with the slow video [AOMI_MOCO_], that was a bit challenging. Maybe because the speeds were much closer as well, so it’s a bit more challenging to imagine, to make sure that I’m not slowing the image to match the video” [participant 12]). In a somewhat contradictory manner, participants who found MI most difficult during AOMI_LOCO_ suggested the scale of the difference between AO and MI processes, and the sheer speed of the imagery, made it hard to accurately generate the imagined *fast-paced* SL-STS (e.g., “I found that one the hardest because it was too distanced between the two [observed and imagined movements] and it was negatively impacting the movements I was imagining due to the mixed signals” [participant 5]).

All participants confirmed that they could successfully engage with MI when instructed to across the experiment. Fourteen participants (66.67%) stated that their MI consisted of only kinesthetic aspects for all conditions (e.g., “there’s [imagining] the initial sort of force production bit, which is really tight on the quadriceps, and then once you’re sort of almost fully extended, that’s when [imagining] the feelings of being balanced comes into it.” [participant 3]). However, seven participants (33.33%) indicated that they sometimes generated visual images as well as kinesthetic images of the SL-STS, with this largely taking place in the MI condition due to a lack of visual stimuli, or in the AOMI conditions when attempting to reset their timing or attention to the task. Of these, three participants (14.29%) imagined seeing her/himself performing the SL-STS from a first-person visual perspective whilst maintaining kinesthetic imagery (e.g., “That [third-person] is the perspective I have of that person doing it [in the video], but not of me doing it, because I’m not on the outside of me, so I imagine seeing and feeling it as if I’m practicing” [participant 7]). Four participants (19.05%) imagined seeing her/himself performing the SL-STS from a third-person alongside their kinesthetic imagery (e.g., “There were probably moments where I was imagining things from a sagittal plane as the video was showing, but I think a lot of the time my focus was on imagining the feeling of and the kind of sensations involved” [participant 16]).

## Discussion

This study aimed to test the DASH (Eaves, Riach, et al., 2016) and VGH (Meers et al., 2020) propositions for *coordinative* AOMI by comparing CSE responses during highly-, moderately-, and lowly-coordinated AOMI for a SL-STS movement. Task-related MI ability and social validation data were also collected to help explain the cognitive processes underpinning any differences in CSE response during these different types of *coordinative* AOMI. CSE was facilitated for AOMI_HICO_ at T3, indicating a combined effect for the observed and imagined *slow-paced* SL-STS. CSE was facilitated for the AOMI_MOCO_ condition at T3 and T2, indicating an effect for the observed *slow-paced* SL-STS and the imagined *medium-paced* SL-STS, respectively. CSE was facilitated for AOMI_LOCO_ at T3 and T1, indicating an effect for the observed *slow-paced* SL-STS and the imagined *fast-paced* SL-STS. However, CSE was facilitated for AOMI_MOCO_ and AOMI_LOCO_ at all three timepoints, AOMI_LOCO_ reported the greatest CSE facilitation at all three timepoints, and facilitation was inversely related to the level of coordination for AOMI at T3 and T2. These findings suggest the increased speed of MI drove the facilitation in CSE across the study as a whole. Overall, the findings lend possible support to both the DASH and VGH propositions for *coordinative* AOMI, as discussed below.

In this experiment, AOMI_HICO_ involved the simultaneous observation and imagery of a *slow-paced* SL-STS movement. Our findings align with both the DASH (Eaves, Riach, et al., 2016) and VGH (Meers et al., 2020) propositions for AOMI_HICO_ as CSE facilitation was increased for this condition at T3, the stimulation timepoint at which the observed and imagined aspects were incongruence for the observed and imagined *slow-paced* SL-STS, compared with BL and AO, but not MI control conditions. This finding agrees with a body of literature showing CSE facilitation for AOMI_HICO_ relative to control and independent AO conditions (e.g., Bruton et al., 2020; Grilc et al., 2024; Wright et al., 2018). Based on the propositions of the DASH (Eaves, Riach, et al., 2016), the CSE facilitation reported for AOMI_HICO_ is caused by the two motor representations for AO and MI of the *slow-paced* SL-STS merging as one sensorimotor stream, leading to widespread activity in the premotor cortex. This merger is likely to cause increased activity in overlapping brain regions for AO and MI, whilst also increasing activity across a wider neural network that involves brain areas solely recruited during AO and MI for the *slow-paced* SL-STS during AOMI_HICO_ (Hardwick et al., 2018; Filimon et al., 2015). An alternative explanation is provided by the VGH (Meers et al., 2020), which argues that the CSE facilitation during AOMI_HICO_ is indicative of a stronger motor representation for MI of the *slow-paced* SL-STS due to the priming effect of AO for the same *slow-paced* SL-STS. The social validation data lends support to this assertion, as participants reported significantly greater MI ability during the AOMI_HICO_ condition compared to all other conditions where MI was instructed in this experiment. Participants reasoned that MI was easier in this condition due to the synchrony between the video demonstration and sonification of the *slow-paced* SL-STS, indicating that the MI component took priority during AOMI_HICO_. In line with the findings of a recent meta-analysis (Chye et al., 2022), CSE facilitation at T3 was descriptively, but not significantly greater for AOMI_HICO_ compared with MI of the *slow-paced* SL-STS. In combination, our CSE and social validation data support the propositions of the VGH that MI is primed during AOMI_HICO_, but indicate that this effect might be minimal, at least for complex whole-body actions like the *slow-paced* SL-STS used in this study.

In the current study, AOMI_MOCO_ involved the observation of a *slow-paced* SL-STS and simultaneous imagination of the feelings and sensations involved with a *medium-paced* SL-STS. Our findings align with the DASH (Eaves, Riach, et al., 2016) proposition for *coordinative* AOMI as CSE was facilitated for the AOMI_MOCO_ condition at T3 compared with all three control conditions indicating an effect for the observed *slow-paced* SL-STS, and at T2 compared with BL and AO control conditions indicating an effect for the imagined *medium-paced* SL-STS. This finding is similar to that reported by Bruton et al. (2020) who showed CSE facilitation for both observed and imagined muscles when controlling for visual attention during *coordinative* AOMI of finger movements, but disagrees with the findings of Meers et al. (2020) and Grilc et al. (2024) that showed CSE facilitation for imagined muscles only during *coordinative* AOMI of finger movements and lower-limb movements, respectively. Based on the propositions of the DASH, the requirement to co-represent two related, but not identical, movements during *coordinative* AOMI involves the maintenance of two parallel sensorimotor representations for the observed *slow-paced* SL-STS and imagined *medium-paced* SL-STS as a set of action affordances. The findings for AOMI_MOCO_ in this study indicate that it is possible to simultaneously co-represent two variations of the same SL-STS movement that differ based on the speed (i.e., *slow-* vs. *medium-paced*) of the simulated movement.

The social validation questionnaire and interview data collected in this study indicated that participants found it difficult to both observe the *slow-paced* and imagine the *medium-paced* SL-STS movements during the AOMI_MOCO_ condition because of the close overlap between the speed of the movements. Based on the propositions of the VGH (Meers et al., 2020), the imagined *medium-paced* SL-STS is prioritized during AOMI_MOCO_, meaning the motor representation is only maintained for this action and the observed *slow-paced* SL-STS serves as a visual guide to facilitate MI processes. If this proposition held true, CSE would only be facilitated at T2 during AOMI_MOCO_, as per recent findings (Grilc et al., 2024; Meers et al., 2020). This was not the case, as CSE facilitation was increased at both T3 and T2 for this condition, a finding that could be explained by attentional mechanisms underlying engagement with AOMI_MOCO_. It is possible that participants directed their attention towards either the observed or imagined SL-STS movements when engaging in AOMI_MOCO_ to account for the cognitive demands of the task, and support the generation and maintenance of imagery as the effortful form of movement simulation. Indeed, eye movement (Bruton et al., 2020) and electroencephalographic data (Eaves, Behmer et al., 2016) shows that individuals switch their attention between observed and imagined actions during *coordinative* AOMI. The interview findings reported here are consistent with this interpretation, suggesting participants may direct more attention to either the observed or imagined action on a trial-by-trial basis, resulting in CSE facilitation at T2 and T3 when aggregated across the AOMI_MOCO_ condition in-full. It is noteworthy that CSE facilitation is increased for AOMI_MOCO_ compared with AO at both T2 and T3, and MI at T3, suggesting an additive effect beyond AO or MI processes in isolation. This is indicative of co-representation of the two related, but not identical, movements during AOMI_MOCO_ and disagrees with the proposition that MI drives the CSE facilitation resulting from *coordinative* AOMI (Grilc et al., 2024; Meers et al., 2020).

In the AOMI_LOCO_ condition used in this study, participants observed a *slow-paced* SL-STS and simultaneously imagined the feelings and sensations involved with a *fast-paced* SL-STS. Our findings disagree with the propositions of both the DASH (Eaves, Riach, et al., 2016) and VGH (Meers et al., 2020) accounts for *coordinative* AOMI as CSE was facilitated for the AOMI_LOCO_ condition at T3 and T1 compared with all other conditions, indicating an effect for both the observed *slow-paced* and imagined *fast-paced* SL-STS during AOMI_LOCO_. This finding contrasts most TMS findings on *coordinative* AOMI, as studies have typically reported increased CSE facilitation for the imagined but not the observed action (Grilc et al., 2024; Meers et al., 2020). Given the low coordination between the movement speeds used for AO and MI components of AOMI_LOCO_, the results also warrant comparison with *conflicting* AOMI. In the only TMS study employing a *conflicting* AOMI condition, Bruton et al. (2020) reported no increase in CSE facilitation in the observed and imagined muscles compared to control conditions (BL, AO). The authors also showed that CSE facilitation was increased in the simultaneously observed and imagined muscle during *congruent* AOMI, and imagined muscle during *coordinative* AOMI, when compared to the *conflicting* AOMI condition. This suggests that the AOMI_LOCO_ condition employed in this study is more representative of *coordinative* than *conflicting* AOMI, but this does not explain the facilitation of CSE for both the observed and imagined SL-STS movements during this condition. It is possible that this reflects the maintenance of two parallel sensorimotor streams for the observed and imagined actions, even when there is a sizable temporal discrepancy between the observed and imagined actions during AOMI_LOCO_, aligning with the sentiments of the DASH (Eaves, Riach et al., 2016).

An alternative explanation, consistent with the VGH (Meers et al., 2020), is that MI is driving the increased CSE facilitation reported for AOMI_LOCO_ at all three stimulation timepoints. The level of force or effort of an imagined task is represented in corticomotor activity, with higher force/effort demands leading to increased CSE facilitation during MI (e.g., Helm et al., 2015; Tatemoto et al., 2017; Tremblay et al., 2001). In this study, CSE increased as imagery speed increased during *coordinative* AOMI for T2 and T3, and the condition that required the participants to imagine performing the fastest SL-STS (i.e., AOMI_LOCO_) had the greatest CSE facilitation across all three stimulation timepoints. It was expected that AOMI_LOCO_ would lead to CSE facilitation at T1, as this is the stimulation timepoint aligned to peak EMG activity for a physically executed *fast-paced* SL-STS. CSE facilitation at T2 and T3 was not expected for this experimental condition. Able imagers demonstrate smaller time differences, known as mental chronometry, between physical performance and MI trials of the same task (Guillot & Collet, 2005). This is logical given the shared neural pathways for movement execution and movement simulation (Hardwick et al., 2018), but more difficult movement tasks typically lead to greater temporal discrepancies between imagined and executed movements (Guillot & Collet, 2005). The social validation data indicated that participants perceived their imagery ability to be lowest during AOMI_LOCO_, with the conflicting speeds for the observed *slow-paced* and imagined *fast-paced* SL-STS movements disrupting their imagery. Despite the use of audio sonification to guide MI during *coordinative* AOMI in this study (Castro, Bryjka, et al., 2021; Castro, Osman, et al., 2021) it is possible that participants timing of the imagery was delayed across some trials for the AOMI_LOCO_ condition due to the complexity of the task, possibly explaining the increased CSE facilitation across all three stimulation timepoints for this form of *coordinative* AOMI.

The findings reported for different forms of *coordinative* AOMI in this study have important implications for motor (re)learning in sports and rehabilitation settings. AOMI_HICO_ is recommended as the optimal form of motor simulation because it addresses the limitations involved with AO (e.g., directs the attention of the learner) or MI (e.g., provides visual movement information to support the generation and maintenance of kinesthetic images) when used alone, and has additive benefits towards motor skill performance compared to these techniques in isolation (Chye et al., 2022; Wright et al., 2022). These performance benefits are supposedly underpinned by synaptic plasticity processes similar to those observed during physical practice (Holmes & Calmels, 2008). The two other types of *coordinative* AOMI employed in this study (AOMI_MOCO_, AOMI_LOCO_) facilitated CSE at stimulation timepoints aligned with the observed and imagined movements. They may therefore have the capacity to benefit (re)learning of variations of the same action or joint actions (Vogt et al., 2013). These *coordinative* AOMI conditions are a viable complementary training method to physical therapy in rehabilitation settings and could be used to promote the (re)learning of actions that are currently impaired or missing from a person’s motor repertoire. For example, a post-stroke patient may benefit from observing videos of themselves accurately performing leg movements such as the SL-STS with their non-affected limb, whilst simultaneously imagining the feelings and sensations associated with performing the same or similar leg movements at different speeds with their impaired limb (e.g., McCormick et al., 2022). In such cases, AOMI_MOCO_ and AOMI_LOCO_ could support motor (re)learning by promoting Hebbian plasticity in a similar manner to that described above for AOMI_HICO_.

All four studies investigating *coordinative* AOMI have used TMS to examine the neurophysiological mechanisms underpinning this form of action simulation (see Bruton et al., 2020; Grilc et al., 2024; Meers et al., 2020; this study), yet behavioral research is lacking. There is a need to empirically test the effectiveness of AOMI_MOCO_ and AOMI_LOCO_ as movement interventions across populations and contexts in order to substantiate propositions that these types of *coordinative* AOMI may be better than AOMI_HICO_ for (re)learning variations of the same movement or joint actions where interpersonal coordination is required (Eaves et al., 2022). Longitudinal research incorporating both neurophysiological and behavioral measures is required to verify the extent to which repeated engagement in different types of *coordinative* AOMI promotes functional connectivity and plasticity changes within the brain, and the association between these neural adaptations and any motor performance and learning improvements after a *coordinative* AOMI intervention period (Bruton et al., 2020; Chye et al., 2022; Grilc et al., 2024).

Single-pulse TMS permits time- and muscle-specific measurement of CSE facilitation (Naish et al., 2014; Rothwell, 1997), and thus allowed the contributions of each simulation state (i.e., the observed and imagined actions) to be distinguished by examining the effects of different types of *coordinative* AOMI on MEP responses in the KE muscle group at three stimulation timepoints. However, this study was limited by only recording MEPs from the lower limb muscles (i.e., the KE muscle group) for the SL-STS during the different *coordinative* AOMI conditions. Future studies should topographically map and record EMG from a wider selection of whole body musculature to draw more conclusive evidence regarding CSE facilitation for different types of *coordinative* AOMI. Additionally, the single-pulse TMS technique adopted in this study only provides an indication of activity associated with *coordinative* AOMI within the motor and premotor cortices of the brain. Activity in other brain regions (e.g., rostral prefrontal cortex; Eaves, Behmer et al., 2016) would not have been represented in the MEP responses recorded in this experiment. There is a need to explore the precise anatomical substrates involved in different types of *coordinative* AOMI using neuroscientific methods with superior spatial resolution to TMS, such as functional magnetic resonance imaging (fMRI). FMRI research employing multi-voxel pattern analysis has shown it is possible to distinguish between different actions for MI and execution (Pilgramm et al., 2016; Zabicki et al., 2016). Applying this analysis to fMRI data for different types of *coordinative* AOMI could further advance the understanding of the neural mechanisms underpinning this form of dual-action simulation and provide more conclusive evidence regarding the propositions of the DASH (Eaves, Riach, et al., 2016) and VGH (Meers et al., 2020) accounts for *coordinative* AOMI.

In conclusion, the findings of this study lend possible support to both the DASH (Eaves, Riach, et al., 2016) and VGH (Meers et al., 2020) propositions for *coordinative* AOMI. Specifically, in partial support of the DASH, all three *coordinative* AOMI conditions resulted in increased CSE facilitation at the stimulation timepoint aligned with the observed SL-STS, and at the respective stimulation timepoints aligned with the imagined SL-STS movements. This aligned with propositions of the DASH that sensorimotor representations for the observed and imagined SL-STS movements would merge during AOMI_HICO_ and AOMI_MOCO_ due to the coordination between the simulated SL-STS movements, but this was not expected for AOMI_LOCO_ because of the increased competition between the observed and imagined SL-STS movements. However, the current study offers more conclusive support for the VGH account of *coordinative* AOMI as CSE was facilitated across all three stimulation timepoints, and CSE increased as imagery speed increased across the *coordinative* AOMI conditions, indicating that imagery is likely driving the CSE facilitation present in the data. However, the study design limits our ability to distinguish between the contributions of AO and MI on a muscle-specific basis during *coordinative* AOMI. This study provides novel neurophysiological evidence supporting for the use of *coordinative AOMI* as an alternative method for (re)learning of movements that extend beyond a learner’s repertoire, or for joint actions that require coordination with the movements of others.

